# Gene expression changes after heat shock of procyclic-form *Trypanosoma brucei* suggest that stress has a role in differentiation to mammalian-infective forms

**DOI:** 10.1101/058701

**Authors:** Igor Minia, Clementine Merce, Monica Terrao, Christine Clayton

**Author notes:** Current address: Max-Delbruck-Centrum für Molekulare Medizin, Robert-Rossle-Str. 10, 13125 Berlin-Buch, Germany (Igor Minia). Current address: Cancer Genome Research Group-NCT, Im Neuenheimer Feld 460, 69120 Heidelberg, Germany.

## Abstract

Trypanosome procyclic forms multiply in the midgut of Tsetse flies, and are routinely cultured at 27°C. Heat shocks of 37°C and above result in general inhibition of translation, and severe heat shock (41°C) results in sequestration of mRNA in granules. The mRNAs that are bound by the zinc-finger protein ZC3H11, including those encoding refolding chaperones, escape heat-induced translation inhibition.a At 27°C, *ZC3H11* mRNA is predominantly present as an untranslated cytosolic messenger ribonucleoprotein particle, but after heat shocks of 37°C - 41°C, the *ZC3H11* mRNA moves into the polysomal fraction. To investigate the scope and specificities of heat-shock translational regulation and granule formation, we analysed the distributions of mRNAs on polysomes at 27°C and after 1 hour at 39°C, and the mRNA content of 41°C heat shocks granules. We found that that mRNAs that bind to ZC3H11 remained in polysomes at 39°C and were protected from sequestration in granules at 41°C. As previously seen for starvation stress granules, the mRNAs that encode ribosomal proteins were excluded from heat-shock granules. Seventy mRNAs moved towards the polysomal fraction after the 39°C heat shock; surprisingly, many of these are also increased when trypanosomes migrate to the Tsetse salivary glands. It therefore seems possible that in the wild, temperature changes due to diurnal variations and periodic intake of warm blood might influence the efficiency with which procyclic forms develop into mammalian-infective forms

**Author summary:** When trypanosomes are inside tsetse flies, they have to cope with temperature variations from below 20°C up to nearly 40°C, due to diurnal variations and periodic intake of warm blood. The procyclic forms, which usually multiply in the midgut, are routinely cultured at 27°C in the laboratory. When they are heated to temperatures of 37°C and above, they shut down protein production, and at 41°C, mRNAs aggregate into granules. We show here that quite a large number of mRNAs are not included in granules and continue to be used for making proteins. Some of the proteins that continue to be made are needed in order to defend the cells against the effects of heat shock. Interestingly, however, a moderate heat shock stimulates expression of genes needed for the parasites to develop further into forms that can colonise the salivary glands. It thus seems possible that in the field, temperature variations might influence the efficiency with which of trypanosomes in tsetse flies become infective for mammals.

## Introduction

Trypanosomes, like all other organisms investigated so far, respond to heat shock by repressing general protein synthesis, while enhancing or retaining synthesis of proteins that are required to survive or recover from heat stress [1]. Unlike other organisms, however, trypanosomes lack the ability to control the transcription of individual protein-coding genes [2–4]. Polymerase II transcription is polycistronic; and monocistronic mRNAs are created by 5’ *trans* splicing of a capped spliced leader (*SL*) and polyadenylation [5]. The selectivity of the heat shock response, like other changes in gene expression, therefore relies on post-transcriptional mechanisms. *Trypanosoma brucei* procyclic forms are the form that grow inside the Tsetse fly midgut. In natural infections, these forms migrate to the proventriculus, developing into epimastigotes, and from there to the salivary glands where they become metacyclic forms which are infective for mammals [6]. Long slender bloodstream forms proliferate in mammalian blood and tissue fluids. Upon reaching high density, they differentiate into non-dividing short stumpy forms [7], which are pre-adapted for differentiation into procyclic forms upon uptake by Tsetse [8].

Nearly all previous work on heat shock in *T. brucei* has concentrated on cultured procyclic forms subjected to a one-hour heat shock at 41°C [1]. Although this is on the upper edge of temperatures that can be tolerated by most Tsetse species in the wild [9], the trypanosomes recover quite rapidly upon return to the normal culture temperature of 27°C [1]. Heating to 41°C inhibits trypanosome transcription initiation [10,11] and stimulates overall mRNA degradation [1], resulting in gradual loss of total mRNA [1]. In addition, translation of most mRNAs is suppressed. After an hour at 41°C, there is almost no mRNA in polysomes, while three types of messenger ribonucleoproten (mRNP) granules appear:these contain most of the mRNA [12] and various combinations of translation factors, the two poly(A) binding proteins PABP1 and PABP2, the helicase DHH1, the aggregation-prone protein SCD6, and the 5’-3’ exoribonuclease XRN1 [1,13,14]. Cycloheximide treatment causes retention of mRNAs in polysomes at 41°C, inhibiting both mRNA degradation and granule formation [1]. Thus as in other organisms, granules are locations for storage and/or degradation of nontranslated mRNAs.

Despite the general shut-down in genome expression after heat shock, synthesis of proteins that are required for survival during, and recovery after, heat shock - such as protein refolding chaperones - continues. We previously showed that the zinc-finger protein ZC3H11 binds to the 3’-UTRs of chaperone mRNAs, and is required both for target mRNA retention and for cellular survival after heat shock [15]. ZC3H11 binds to MKT1 and to PBP1, which in turn recruits LSM12 and poly(A) binding proteins PABP1 and PABP2 [16]. MKT1 and PBP1 remain distributed throughout the cytosol after heat shock, whereas they colocalise with SCD6 in the stress granules that form after starvation [16].

Recently, we investigated how ZC3H11 itself is regulated [12]. ZC3H11 is barely detectable in both bloodstream and procyclic forms grown at their normal culture temperatures of 37°C and 27°C respectively [15]. When procyclic forms are incubated at 37 - 41°C, the level of ZC3H11 protein progressively increases. This is partly caused by a loss of protein degradation, but more prominently by translational control. At 27°C, the *ZC3H11* mRNA migrates in sucrose gradients as a messenger ribonucleoprotein particle at or just above the small ribosomal subunits, but after a 1h heat shock at 37°C, 39°C or 41°C, nearly all of the *ZC3H11* mRNA is in the polysomal fractions and *ZC3H11* mRNA does not colocalise with heat shock granules [12]. In this paper we have examined whether other mRNAs show similar translation regulation after a 39°C heat shock, and identified additional mRNAs that escape granule formation after a 41°C heat shock.

## Methods

### Trypanosome culture

Trypanosome culture conditions were as described in [17]. Procyclic trypanosomes were grown in MEM-Pros medium at 27°C (unless stated otherwise) at densities lower than 6×10^6^ cells/ml. All experiments were done with Lister 427 monomorphic procyclic form parasites expressing the *Tet* repressor.

### Polysome analysis and RNASeq

3-5× 10^8^ procyclic cells were treated with cycloheximide (100μg/ml) for 5minutes, harvested at room temperature by centrifugation (850g, 8min, 20°C), washed once in 1ml of ice-cold PBS and lysed in 300μl of lysis buffer (20mM Tris pH7.5, 20mM KCl, 2mM MgCl_2_, 1mM DTT, 1200u RNasin (Promega), 10μg/ml leupeptin, 100μg/ml cycloheximide, 0.2% (vol/vol) IGEPAL) by passing 20-30 times through a 21G needle. After pelleting insoluble debris by centrifugation (17000g, 10min, 4°C) and adjusting to 120mM KCl, the clarified lysate was layered onto a 17.5-50% sucrose gradient (4ml) and centrifuged at 4°C for 2 hours at 40000 rpm in Beckman SW60 rotor. Monitoring of absorbance profiles at 254nm and gradients fractionation was done with a Teledyne Isco Foxy Jr. system. A human β-globin *in vitro* transcript was sometimes added to each of the collected fractions as a spike-in control. RNAs from pooled fractions were purified using Trifast.

### Protein characterisation

Proteins were detected by Western blotting according to standard protocols. For detection of the endogenous ZC3H11 protein only cytoskeleton-free extracts were used. Antibodies used were to the ZC3H11 (rabbit, 1:10000, [12]), RBP6 [18] and PTP1 [19]. Detection was done using ECL solutions (GE Healthcare).

### Purification of trypanosome heat shock granules

Granules from normal and heat-shocked procyclic cells were enriched as described previously [20]. 5×10^8^ control or heat-shocked (1 hour at 41°C) procyclic cells were harvested at room temperature by centrifugation (1500g, 10min), washed in 1ml of PBS and lysed in 200μl of ice-cold buffer A (20mM Tris-HCl pH 7.6, 2mM MgCl_2_; 0.25M sucrose, 1mM DTT, 10% glycerol, 1% TritonX100, 800u RNasin (Promega), 1 tablet Complete Protease InhibitorCocktail EDTA free (Roche)/10ml buffer) by pipetting. Lysis was confirmed microscopically. The lysate was clarified (20000g, 10min) and the supernatant (SN1) was transferred to fresh tube with 750μl of peqGOLDTrifast FL(Peqlab). All remaining supernatant was removed after one short centrifugation (3min, 20.000g). The pellet was resuspended again in 200μl of buffer A by passing 30-40 times through a 21G syringe, vortexed and centrifuged (20000g, 5min). The supernatant (SN2) was taken and the pellet was resuspended in 200μl buffer A as above. The whole procedure was repeated one more time to obtain the supernatant SN3. Then the pellet was resuspended one more time in 200μl buffer A as above and microtubules were disrupted by the addition of 12 μl 5M NaCl (283mM final conc.), the samples were passed through 21G syringe, incubated on ice for 30 minutes with vortexing every 5 minutes, then centrifuged (20000g, 10min). The supernatant (SG) was removed and used to prepare the "small granule" RNA (SG). The pellet was washed once in 200μl of buffer A without resuspension (20000g, 10min) and finally resuspended in 750μl of Trifast FLto make the "large granule" (LG) RNA. Another 5×10^7^ control or heat-shocked procyclic cells were taken to obtain total RNA.

### Sequencing and sequence analysis

Total RNA was incubated with oligonucleotides complementary to trypanosome rRNA and RNAse H, and integrity was checked by Northern blotting woth a probe that detectes the beta-tubulin mRNA. The samples were then subjected to high throughput sequencing such that most samples gave about 30 million aligned reads. Sequences were aligned to the latest available *T. brucei* TREU927 genome sequence using Bowtie [21], allowing for up to 20 sequence matches. Reads that aligned to open reading frames were then aligned using a custom script, again allowing for each read to align up to 20 times. To extract the reads for individual open frames, we used a modified version of the "unique open reading frame" list of Siegel et al. [22]. Reads per million and other routine calculations were done in Microsoft Excel. Differences in RNA abundance between conditions or fractions were assessed using DeSeq [23]. Untranslated region sequences were downloaded from tritrypDB and sequence motifs searched using DREME and MEME [24]. Other statistical analyses were done in R. Functional gene classes were assigned manually using a combination of automated annotations and publications. All raw sequence data are available at Array Express with accession numbers E-MTAB-4555 (polysomes) and E-MTAB-4557 (granules).

## Results

### Heat shock increases polysomal loading of a subset of mRNAs

The first part of our study concerned the movement of mRNAs into, and out of, the polysomal fraction after a one-hour heat shock at 39°C. We were particularly interested in knowing which mRNAs show regulation similar to that of ZC3H11, since we hoped in that way to identify conserved sequence motifs. We chose 39°C because preliminary results showed that the treatment was sufficient to move *ZC3H11* mRNA into the polysomal fraction, while only partially inhibiting overall translation. It is also a treatment that could be tolerated by Tsetse flies [9]. Lysates from procyclic-form trypanosomes with or without heat shock were fractionated on sucrose gradients, which were then divided into free, subunit, light polysome and heavy polysome fractions (Fig 1A). The 39°C heat shock caused a shift of the ribosomes from the polysomal towards the free subunit and monosome fractions (Fig 1A). To find the proportion of mRNA that was in each fraction, we analysed samples by Northern blotting, using the spliced leader as probe (Fig 1B) and including inputs (non-fractionated samples) as controls. The total amount of mRNA from the 39°C-treated cells was 57% of that from the non-shocked samples, but the sucrose gradient distribution of mRNA was similar to that of the non-shocked parasites (Fig 1B and S1 Table, sheet 2). This suggests that loss of translation is associated with mRNA degradation. (We cannot tell which comes first). All samples were subjected to RNASeq (S1 Table, sheet 3, and S1 Fig).

**Fig. 1.**
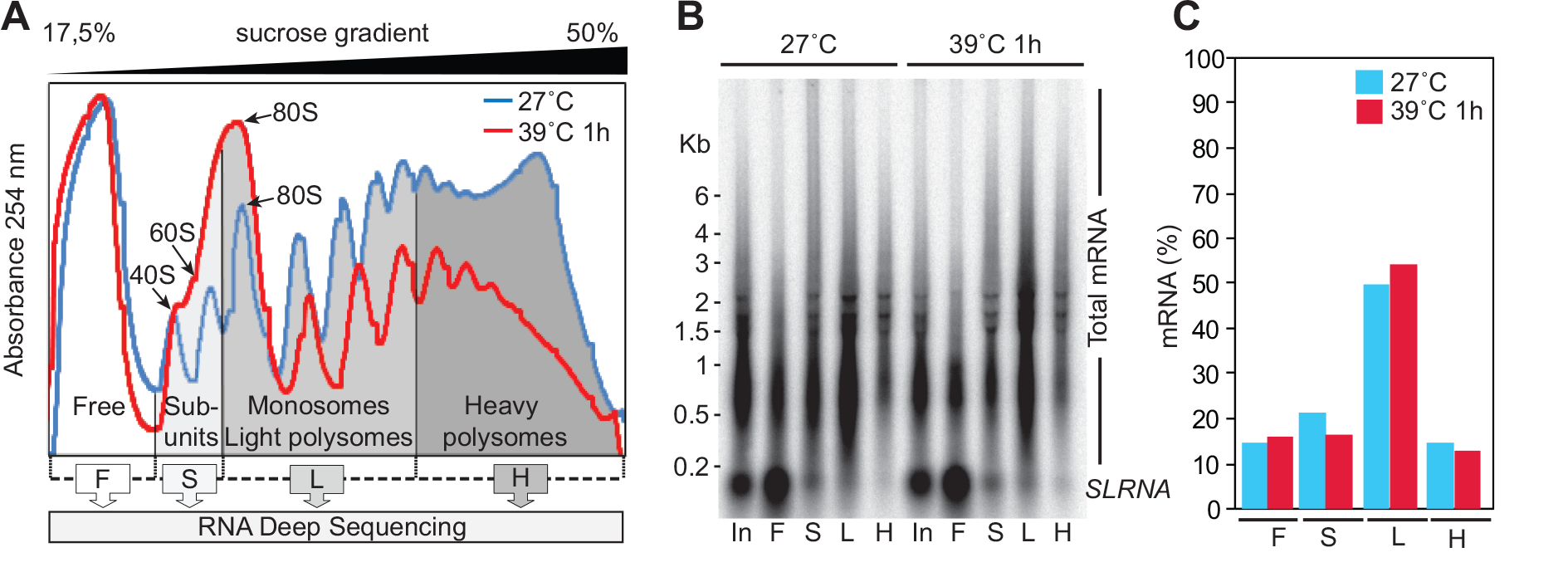
Transcriptome-wide polysome profiling upon heat shock of procyclic forms A. Representative polysome profiling of procyclic cells grown at 27°C (normal culture conditions) or heat-shocked for 1h at 39°C. Fractions were pooled as described and used for the analysis (F:free, S:ribosomal subunits, L:monosomes and light polysomes, H:heavy polysomes). B. For normalization, RNA prepared from four pools was analysed by Northern blotting (NB). The total signal from spliced mRNA (present at the 5’-end of each trypanosomal mRNA) was used to calculate the distribution of total mRNA in four pools (mean, n=2). C. Changes in translation of individual mRNAs after heat shock. The ribosome load was determined for each detectable TriTrypDB-annotated mRNA (2 biological replicates).

To find out the proportions of each mRNA in the sucrose gradient fractions, we normalised the read counts / million reads (S1 Table, sheet 3) according to the spliced leader signals (S1 Table, sheets 4 and 5). We then calculated the percentage of each mRNA that was in the different sucrose gradient fractions (S1 Table, sheet 6). (A summary of the control results at 27°C was included in [4].) mRNPs migrate in the monosome or free fractions. In the following discussion we will assume that mRNAs that migrate in the denser part of the gradient are being actively translated. However, although there are no microscopically visible granules at 39°C, some association of mRNAs with smaller aggregates cannot be ruled out. The correlation coefficients for percentage in polysomes between replicates ranged from 87% to 99% (S1 Fig). On average, 50-70% of the transcript reads were in the polysomal fractions (Fig 2A "All"), and there was a statistically significant (but very small) increase in the percentage of the mRNA that was polysomal after heat shock.

**Fig. 2.**
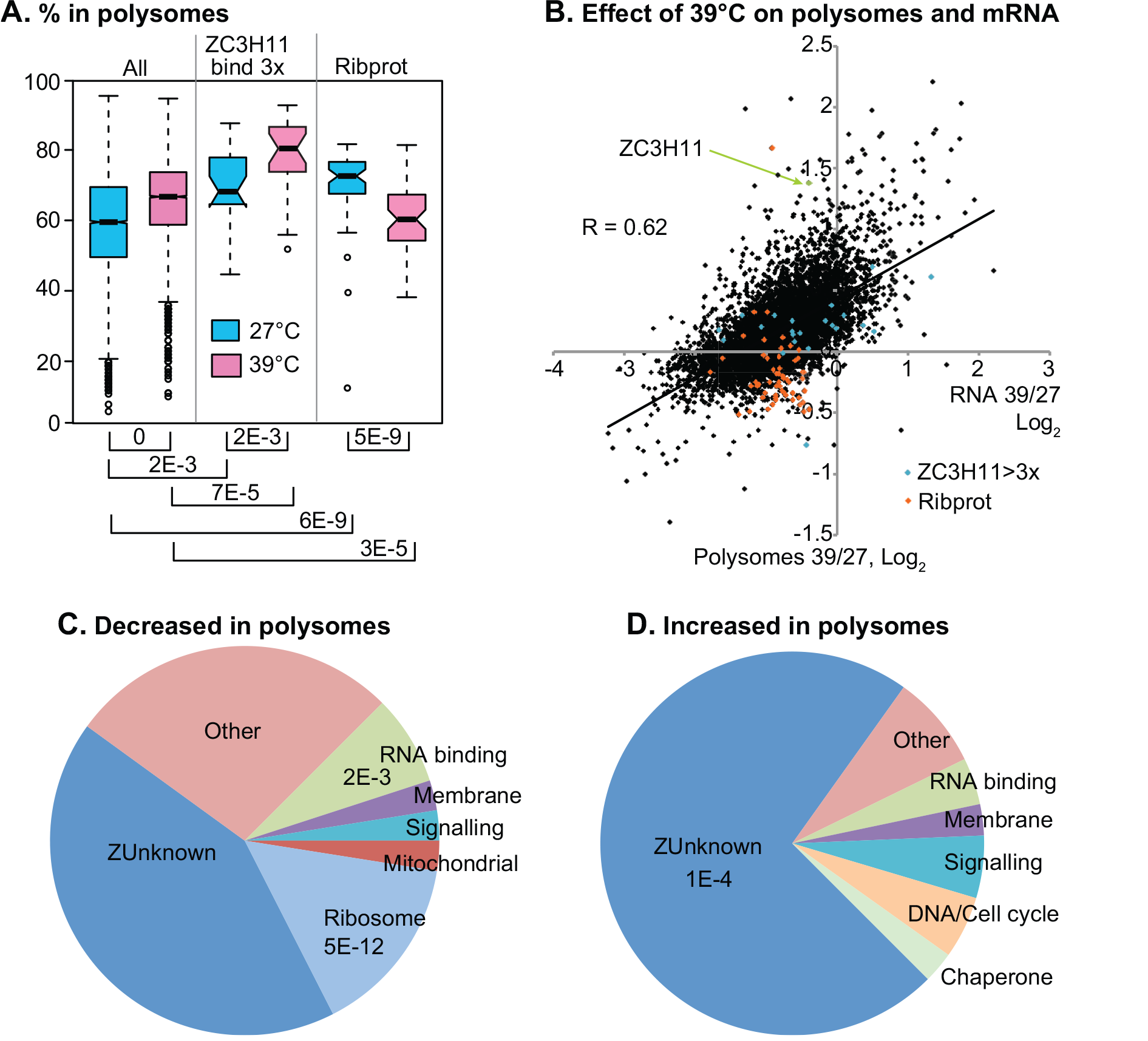
Analysis of mRNAs in polysome gradients (A) Genes were divided into three categories:"Decrease" means that the proportion of mRNA in polysomes (heavy and light combined) decreased by 1.25x or more after heat shock; "Increase" means that the proportion in polysomes increased by 2x or more; and "Other" are the remaining genes. Results are displayed as box plots (25th to 75 percentiles with central median). Notches represent the 95% confidence intervals for the medians. Dotted lines show 1.5x the inter-quartile range and circles are outliers. (B) Correlation between polysome loading and abundance. n each case we show the log2 of the value for 39°C divided by the value for 27°C. The X-axis is the amount of total RNA and the y axis is the percentage in the polysomes (heavy and light combined). (C) Functional classes of mRNAs that moved to polysomes after heat shock. The results of a Fisher test for significiant enrichment are shown. (D) Functional classes of mRNAs that shifted away from polysomes after heat shock.

There were 80 transcripts for which the percentage in polysomes decreased by a factor of 1.25 or more after heat shock (S1 Table, sheet 8). For this group, the percentage on polysomes was higher than average at 27°C, and lower than average after an hour at 39°C (Fig 2A). A subset of these mRNAs was distinguished by poor translation even before heat shock (S2 Fig, subset A):it includes *RBP33*, cis-spliced poly(A) polymerase, *PAG2* and *PAG4* mRNAs. Transcripts encoding ribosomal proteins were notably enriched in the set of mRNAs with decreased translation (Fig 2A-C, S1 Table, sheet 9):

77 mRNAs, including that encoding ZC3H11, had a two-fold higher proportion in polysomal fraction after heat shock (S1 Table, sheet 7). This group had almost universally been in the lighter fractions at 27°C. It was enriched in mRNAs encoding proteins of no known function (Fig 2C, S1 Table sheet 9). More detailed analysis of these mRNAs placed them into three categories (S3 Fig). Most of the mRNAs migrated in the free fraction, lighter than the monosomes, in the 27°C samples. The exception was group (B), which included *ZC3H11*. These mRNAs were distinguished from the others by the fact that the mRNAs migrated mainly with monosomes at 27°C. We have shown for *ZC3H11* that this is not due to association with a small ribosomal subunit [12] and the reason for the different behaviour is unknown. Group (A) showed weaker movement towards polysomal fractions after heat shock than the other groups. The mRNAs that can bind to ZC3H11 showed slightly higher than average polysome loading at both 27°C and 39°C (Fig 2A).

To find changes in overall abundances of total and polysomal mRNAs, we compared the total counts (S2 Table, sheets 1 and 2). 260 mRNAs were significantly (>2x, Padj<0.01) increased in relative abundance after heat shock (S2 Table, sheet 3). However, since the total amount of mRNA had decreased by about 40% after the heat shock, the numbers of copies per cell of most mRNAs were reduced (Fig 2B). Those mRNAs for which polysomal association increased tended to show less severe degradation after heat shock (Fig 2B). Again it is not possible to assign cause and effect since translation could influence degradation and *vice-versa.*

The mRNAs that are able to bind ZC3H11 encode proteins that are always needed in high amounts, so these mRNAs were correspondingly strongly polysome associated at 27°C (Fig 6A). This is probably ZC3H11-independent:ZC3H11 is barely detectable at this temperature and RNAi has no effect on cell proliferation or morphology [15]. Association of ZC3H11 targets with polysomes was more marked at 39°C, when ZC3H11 is expressed (Fig 6A), but the relative increase was not significantly different from that of the bulk mRNA population (Fig 6B).

### Heat shock at 39°C changes the abundances of mRNAs associated with differentiation

As expected, the mRNAs whose relative abundances increased at least 2-fold after one hour at 39°C, or which moved from non-translated to polysomal fractions, included mRNAs encoding several chaperones, including two ZC3H11 targets (Tb927.10.16100 and Tb927.2.5980) (Table1). There was also a moderate increase in the major cytosolic HSP70 mRNA. The surprise came when we compared this group of mRNAs with transcriptomes from various developmental stages:we found a very significant overlap with mRNAs that are increased during the differentiation of procyclic-form trypanosomes to epimastigotes, metacyclic forms, and bloodstream forms (Fig 3A & B and S2 Table, Sheet 3). Even three of the chaperones were in this category. Notable among the epimastigote-or salivary-gland-specific genes were several that are associated with meiosis, MND1, HOP1, SPO11 and MSH5 (Table1). The MND1 homologue mRNA Tb927.11.5670 was not only increased in the total RNA, but also moved towards polysomes (45% in polysomes at 27°C, 74% at 39°C). The HOP1 homologue mRNA Tb927.10.5490 showed a similar shift towards the polysomal fraction, but no RNA abundance change. YFP-tagged versions of both proteins are restricted to the nuclei of epimastigote-like cells in salivary glands [25]. SPO11 is probably also meiosis specific. MSH5 is annotated as a putative meiosis mis-match repair protein but there is no experimental evidence for this and the assignment is based on rather ambiguous BLAST matches. Apart from these, mRNAs encoding several putative cell cycle regulators and a telomere-binding protein were increased in abundance or polysome association (Table1).

Examination of mRNAs that decreased in abundance showed that they ere spread over numerous functional categories. These mRNAs significantly overlapped with mRNAs that decrease during differentiation of procyclic forms to epimastigotes or bloodstream forms (Fig 3C). There was in contrast no significant overlap with mRNAs that decrease in stumpy bloodstream forms [26].

**Fig. 3.**
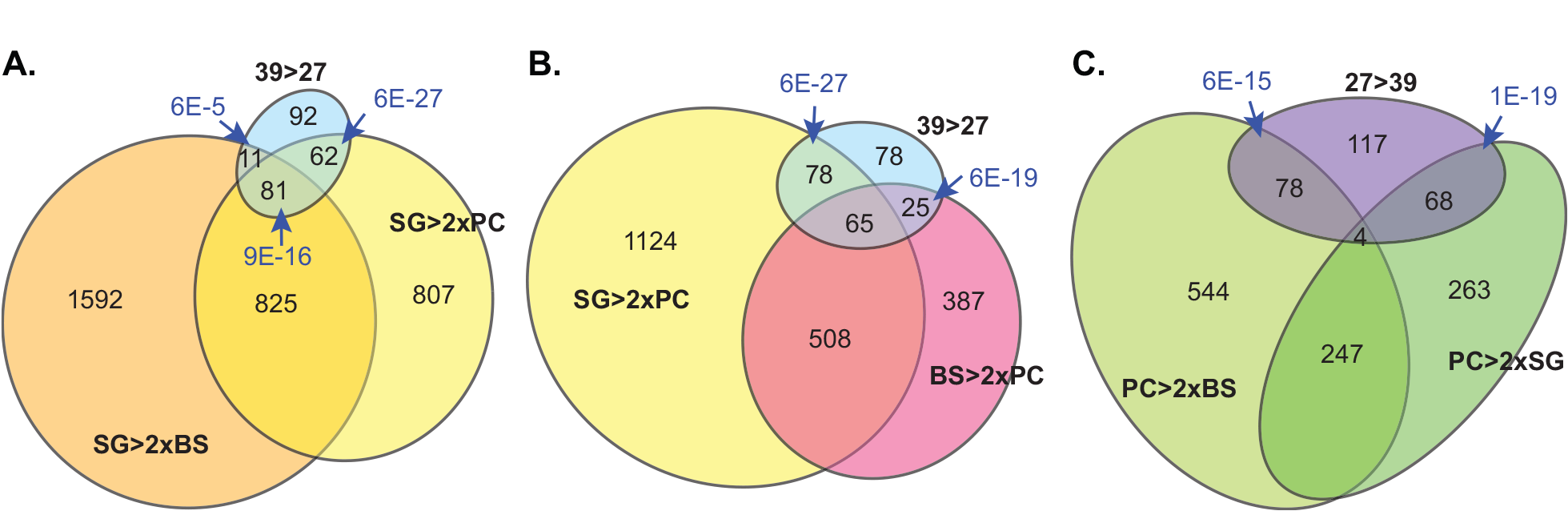
Heat shock induces developmentally regulated mRNAs (A) The mRNAs that increased in abundance after heat shock (Padj <0.01 and 39/27 >2) were compared with mRNAs that are >2x more abundant in slaivary gland (SG) trypanosomes than in procyclic form (PC) trypanosomes, and with mRNAs that are >2x more abundant in salivary gland (SG) trypanosomes than in bloodstream form (BS) trypanosomes. The overlap between the last two categories includes over 900 mRNAs that are most abundant in the salivary gland trypanosome population. The probability that the overlaps would arise by chance (Fisher test) are indicated using blue arrows. Each test was done for a complete overlap, i.e 39>27 overlap with SG>2xBS and 39>27 overlap with SG>2xPC, as well as for the overlap between all three categories. (B) The mRNAs that increased in abundance after heat shock were compared with mRNAs that are >2x more abundant in salivary gland (SG) trypanosomes than in procyclic form (PC) trypanosomes, and with mRNAs that are >2x more abundant in bloodstream form (BS) trypanosomes than in procyclic form (PC) trypanosomes. Fischer tests are for the two-class overlaps. (C) The mRNAs that decreased in abundance after heat shock (Padj <0.01 and 39/27 <0.5) were compared with mRNAs that are >2x less abundant in salivary gland (SG) trpanosomes than in procyclic form (PC) trypanosomes, and with mRNAs that are >2x less abundant in bloodstream form (BS) trpanosomes than in procyclic form (PC) trypanosomes.

**Table 1.**
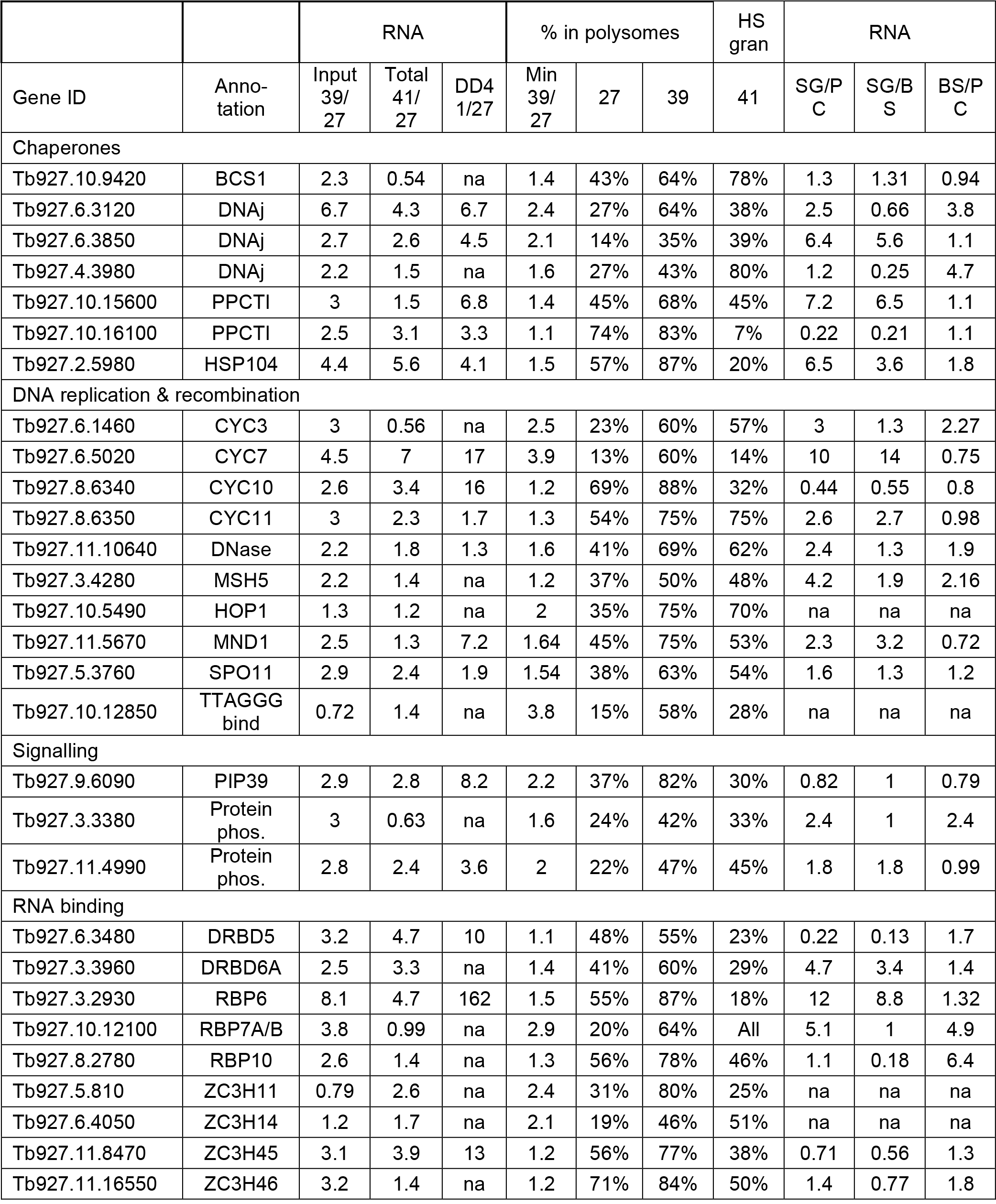
Selected mRNAs that show increased translation or abundance at 39°C. For increased abundance the threshold was a 2-fold relative increase with adjusted P value of less than 01 in DESeq. The ratio for input has not been corrected for the mRNA content of heat-shocked cells (57% relative to 27°C). Other mRNAs that showed a minimum (min) increase of 2-fold in the percentage of that RNA that was in polysomes. RNA abundance ratios for Droll et al. [15] are indicated by "DD". salivary glands (SG) are taken from [27] and ratios for bloodstream form (BS) versus procyclic form (PC) are from [28]. HS gran:percentage in heat shock granules at 41°C. na:no data available. "PPCTI" = peptidyl-prolyl cis-trans isomerase; "Protein phos" = protein with protein phosphatase domain. Ratios are shown to 2 significant figures. A complete list of ORFs that were regulated at the level of total RNA in at least 2 experiments is in S2 Table, sheet 11.

### Regulators of differentiation are induced at 39°C

The numerous changes in developmentally regulated mRNAs after an hour at 39°C suggested that some regulatory proteins might also have been affected. Indeed, mRNAs encoding 9 potential RNA-binding proteins were conserved increased after heat shock (Table 1). Of these, two-*DRBD6* and *RBP6* - peak in salivary gland parasites [27]. Only 55% of the *RBP6* mRNA was in polysomes at 27°C, but 87% was in the fraction after heat shock. Induced expression of RBP6 in procyclic forms is known to trigger the the procyclic-epimastigote-metacyclic differentiation cascade [18]. *RBP10* mRNA is most abundant in growing bloodstream forms [29]. Expression of ZC3H14 and ZC3H45 proteins has not yet been detected but the ZC3H45 mRNA is preferentially translated in bloodstream forms [30]; and ZC3H46 protein is more abundant in bloodstream forms than procyclics [31]. RBP7 protein is present in slender bloodstream forms and increased in stumpy forms [32,33]. RNAi targeting RBP7 increases prevents cAMP -induced stumpy-form differentiation, while overexpression of RBP7 causes G_1_/G_0_ cell cycle arrest and causes initial gene expression changes associated with procyclic differentiation [34].

In addition to these RNA-binding proteins, heat shock induced mRNAs encoding three protein phosphatases. Two of these have no known function, but *PIP39* mRNA is higher in stumpy and procyclic forms than in bloodstream forms, and PIP39 becomes phosphorylated during stumpy-to procyclic differentiation [35]. Movement of *PIP39*, *RBP6* and the Tb927.11.4990 kinetoplastid-specific phosphatase mRNA to polysomes, as well as some others with a similar pattern, was confirmed by Northern blotting (Fig 4A, B). Finally, to see whether a more moderate heat shock might also trigger changes in differentiation regulators, we grew procyclic forms at 37°C overnight. Indeed, the levels of both RBP6 and PIP39 proteins were increased (Fig 4C).

**Fig. 4.**
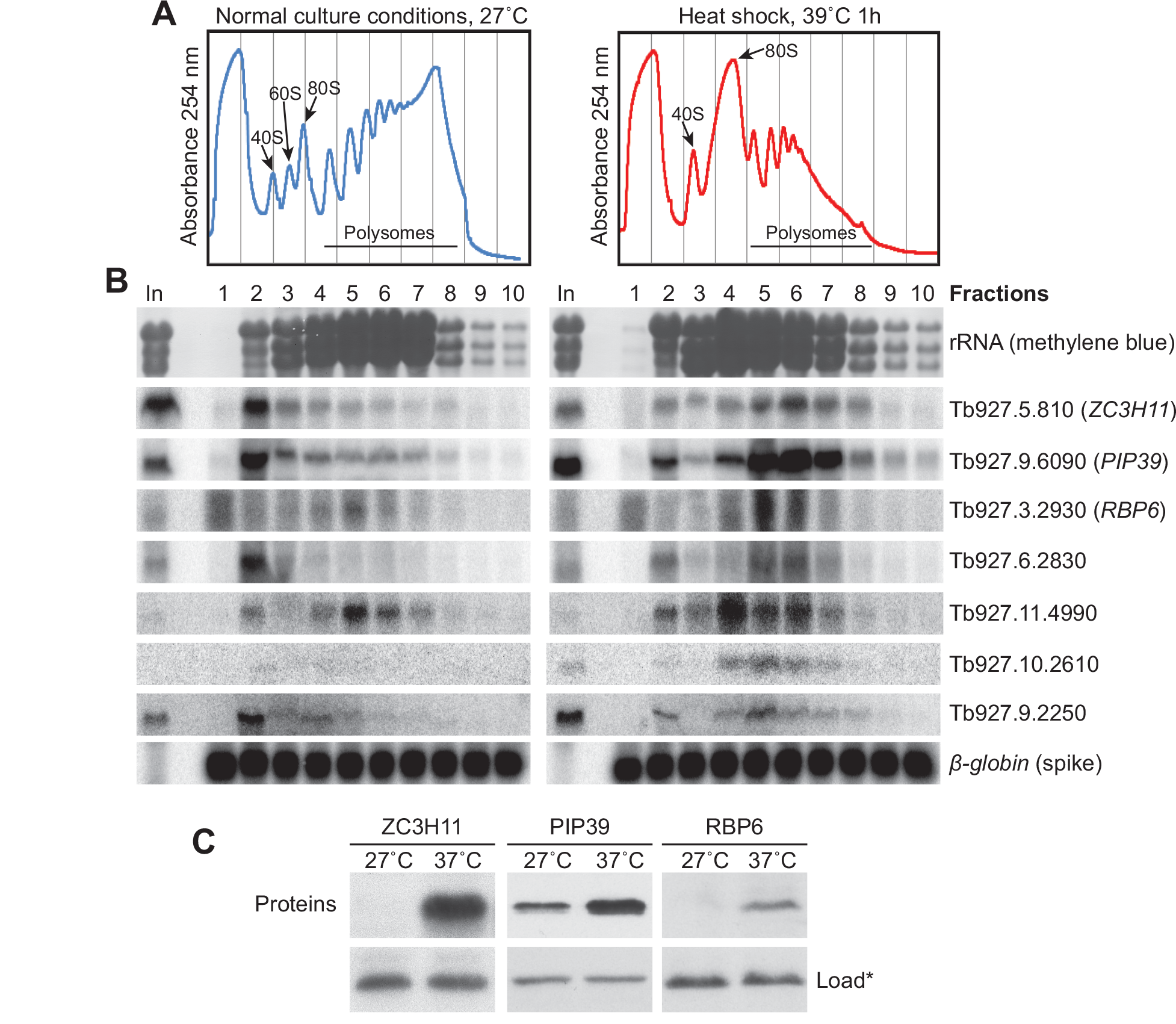
Confirmation of results for individual mRNAs. (A) Extracts from cells grown ta 7°C, or subjected to a 39°C heat shock, were separated on sucrose gradients:some typical gradients are shown. (B) Northern blots of chosen mRNAs that moved from the non-polysomal to the polysomal fraction. A typical rRNA profile is shown as a control. In vitro transcribed human β-globin RNA was added as a spike-in to each fraction before RNA preparation, and is shown as a control of equal RNA isolation efficiency. C. ZC3H11, PIP39 and RBP6 protein levels before and after chronic mild heat shock (16h at 37°C) analyzed by Western blotting. An unspecific band (*) recognized by the anti-ZC3H11 antibody is shown as loading control.

### An AU repeat in the 3’-untranslated region of the ZC3H11 mRNA increases expression independent of heat shock

*ZC3H11* translational control is specified by part of the*ZC3H11* 3’-untranslated region (3’-UTR) [12]. We wondered whether the mRNAs with increased polysomal localization after heat shock had any motifs in common with *ZC3H11* mRNA. We therefore compared the 3’-UTRs of these mRNAs with a control group of 3574 mRNAs that had decreased polysomal loading (39°C/27°C less than 0.7). We found no enriched motifs. We then tried subdividing the heat-induced mRNAs. First we examined 31 3’-UTR sequences from mRNAs that were not only moved towards polysomes, but also 2x increased in abundance in salivary glands [27]. 21 of these had a (U/A)UUAUUUU motif, compared with 731 controls (E-value 6e-4); the motif is present four times in the ZC3H14 mRNA. Seven of the 8 mRNAs showing close co-regulation with *ZC3H11* (group A in S3 Fig) shared the sequence ACT(T/G)CCC (E=2.5e-4) and all eight had A(A/G)TGAAA (E=0.01). Both sequences are found within the 524-nt regulatory segment of the *ZC3H11* 3’-UTR, but not in the core essential regulatory region of 71nt [12]. No shared enriched motifs were found in group B from S3 Fig.

The 71nt *ZC3H11* core regulatory region functioned only in the context of sequences from nt 1-645 of the 3’-UTR [12]. Using a plasmid reporter, we found that an AU repeat at the beginning of this region (the start of the *ZC3H11* 3’-UTR) increased expression 3-fold independent of heat shock (Fig 5A, B). To see whether AU repeats were generally associated with high translation, we divided mRNAs with annotated 3’-UTRs into two categories:those with at least 9 contiguous (AU) dinucleotides and those without. Relevant gene lists are in S3 Table. The mRNAs with at least one copy of (AU)_9_ were slightly, but significantly better translated, as judged by both the proportion of the mRNA in polysomes (S1 Table) and the ribosome density [4] (Fig 5C). They also had slightly longer half-lives [28] (Fig 5C). The presence of 9 or more (AU) in tandem therefore seems to enhance mRNA abundance and translation, but has nothing to do with heat shock regulation.

**Fig. 5.**
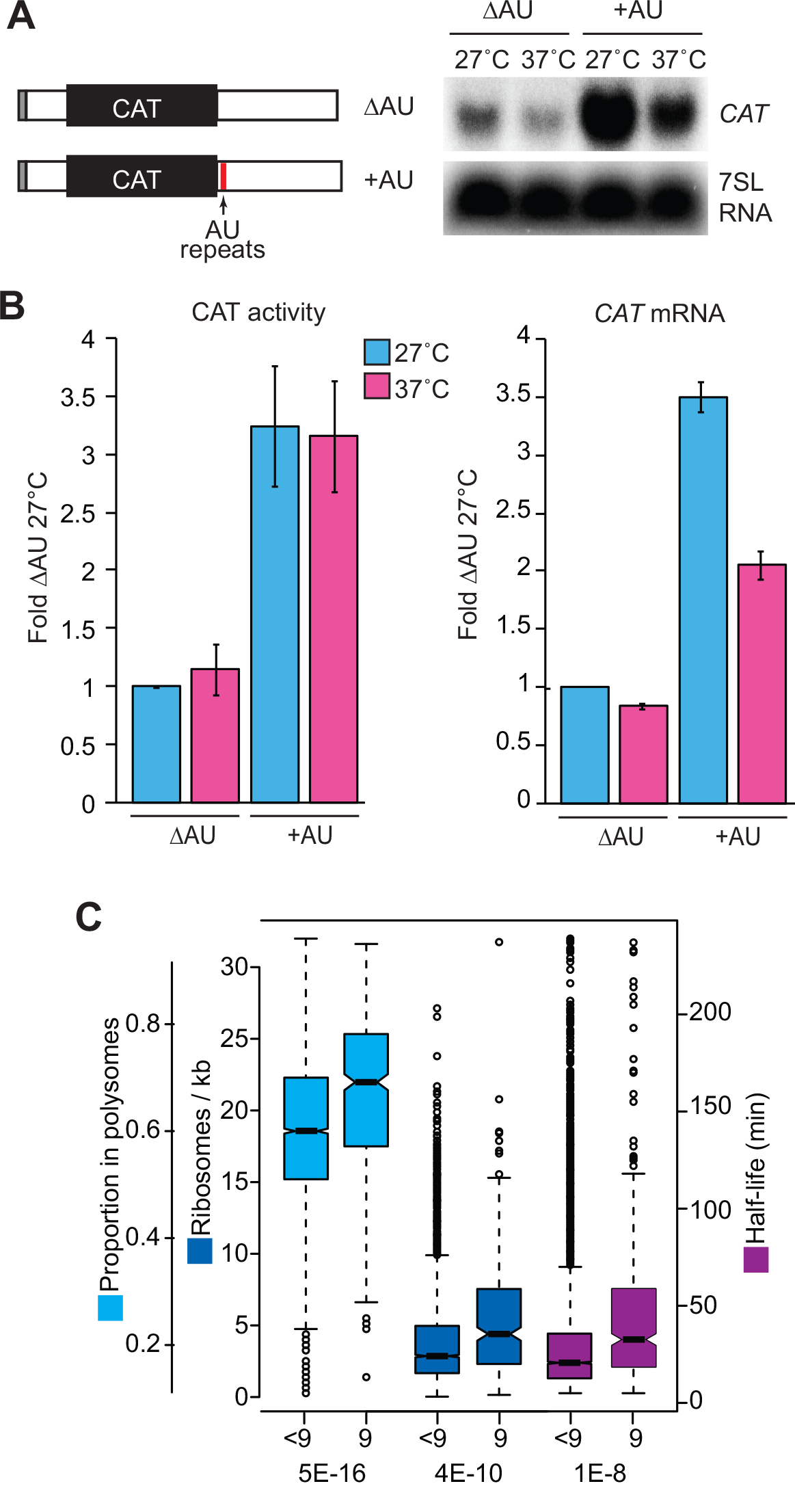
A poly(AU) motif enhances expression. A. *CAT* reporter mRNAs with or without poly(AU) repeats within a truncated ZC3H11 3’-UTR (nt 1-645) [12] were detected by Northern blotting (NB), using cells with or without chronic mild heat shock (16h at 37°C). B. Relative *CAT* mRNA and CAT activity levels at 27°C and after the heat shock (A) were determined by Northern blotting and CAT assay (mean ± SD, n=3). C. mRNAs with at least one copy of (AU)9 (”9”) were compared with mRNAs with less than 9 contiguous copies of (AU) with regard to proportion in polysomes (this paper), ribosome density [4] and mRNA half-life [28]. Results are displayed as box plots (25th to 75 percentiles with central median). Notches represent the 95% confidence intervals for the medians. Dotted lines show 1.5x the inter-quartile range and circles are outliers. For each set of values, the probability that there is no difference between the two sets (student t-test) is shown beneath the relevant columns. For half-lives, only mRNAs with values between 5 min and 240 min were considered.

### At 27°C, less than 5% of mRNAs are in structures with diameters exceeding 24nm

Our second series of experiments addressed mRNA targeting to heat shock granules, which form only at temperatures of at least 40°C [1]. Lysis of trypanosomes in the absence of detergent results in trapping of structures with a diameter of more than 24nm within the microtubule corset [20]. (For comparison, a ribosome is just under 3nm across.) The trapped material can then be released with detergent and fractionated by by ultracentrifugation, which yields a small granule (SG) supernatant and a large granule (LG) pellet. First, we examined cells growing at 27°C. The SG fraction contained about 3.2% of the mRNA, and the LG pellet just 0.8%, as judged by hybridisation with a spliced leader probe [12] (S4 Table, sheet 2). We subjected duplicate fractions to RNASeq (S4 Table, sheets 3 and 4). To work out the proportion of each mRNA within the SG and LG fractions, we compared those results with those for total RNA (S4 Table, sheet 3). The replicates for total RNA of cells growing at 27°C did not correlate very well (S4 Fig A), perhaps because the cells had somewhat different cell densities at the time of harvest (about 3.7 × 10^6^ and 5 × 10^6^/ml). For the 27*C samples we therefore also compared the granule results for individual replicates (S4 Table, sheet 4) with those for the input in the polysome experiments (S4 Table, sheet 5, S4 Fig A). Independent of the way the calculation was done, the proportion of mRNA that was trapped inside the microtubule corset in normally growing cells was determined mainly by the length of either the open reading frame or the complete mRNA (Fig 6A and S4 Fig A, B). This suggests that the trapping was simply due to the size of the polysome and had nothing to do with regulation or granule formation. There was no significant correlation between the percentage in granule fractions and the percentage in polysomes at 27°C (Fig 6B). It was however notable that mRNAs encoding ribosomal proteins were not trapped in granule fractions at all. Even allowing for the short length of most ribosomal protein mRNAs (Fig 6A), their behaviour was anomalous (S5 Table).

**Fig. 6.**
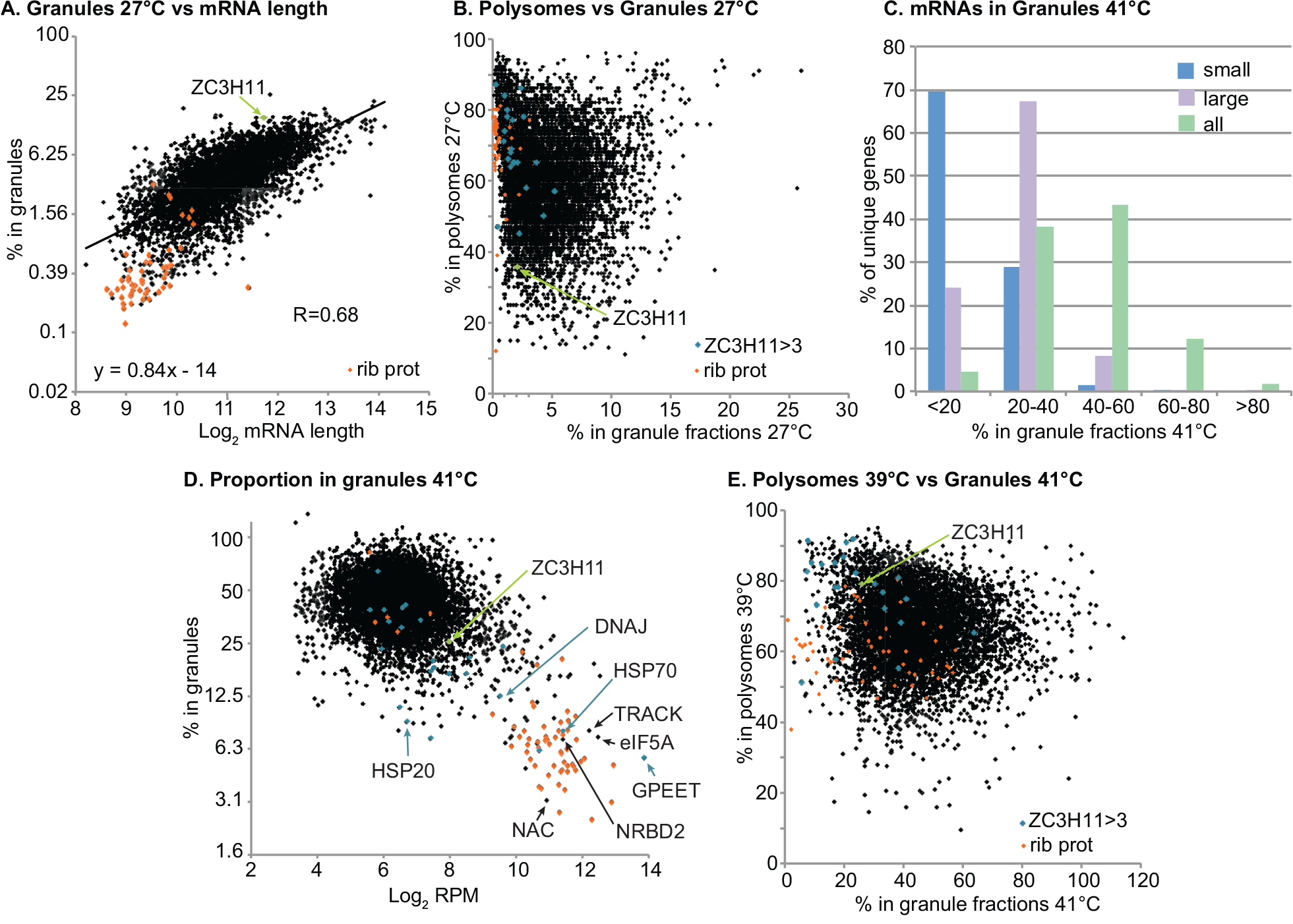
Granule formation at 27°C and 41°C. (A) The percentage of mRNAs in the small granule fraction was plotted against coding sequence (CDS) length. Both axes are on a log2 scale and the correlation coefficient was calculated using log-transformed values. (B) Box plots (as in Fig 5) showing the relationship between the percent of mRNAs in all granules (small and large combined) and the percent in polysomal fractions. Genes encoding ribosomal proteins were treated separately. The number of CDSs in each group are indicated below. The T-test results at the bottom compare each result with a single control:for 27°C, this is the group of mRNAs with less than 2% in the small granule fraction, and for 41*C it is those with less than 20% in granules. T-test results between neighbouring sets are shown above the boxes.

### Binding to ZC3H11 correlates with protection from heat shock granule recruitment

We next examined the effect of a 41°C heat shock. First, we compared results for total mRNAs with those obtained at 39°C, and also with previously published results. The variability in the 27°C dataset meant that P-values for the total RNAs were high (S5 Table) and the overall correlation between different experiments was poor (S1 Fig). This probably reflects differences in cell density as well as temperature. However, a core set of mRNAs was increased in at least 2, and often all three, datasets (Table 1 and S2 Table, sheet 11). In addition to a few chaperone mRNAs, these once again included mRNAs indicative of developmental regulation:CYC7, CYC11 and CYC10; SPO11 and MND1; bloodstream-specific alternative oxidase, pyruvate kinase and GPI-PLC; 6 protein kinases; 3 protein phosphatases including PTP1; and 8 RNA-binding proteins including both RBP10 and RBP6. After one hour at 41°C, 6% of the total mRNA was in the small granule fraction, and 19% in large granules (S4 Table, Sheet 2). At the level of mRNAs from individual genes, however, the distribution looked very different. This is because half of the sequence reads were contributed by the most abundant 10% of the transcripts. For most coding sequences, 20-60% of the mRNA was in one of the granule fractions, usually with the large granule fraction predominating (Fig 6C, D). In contrast, a subset of very abundant mRNAs was not associated with granules (Fig 6D). These included those encoding ribosomal proteins (Fig 6D, Fig 7A and S4 Fig C,D), procyclin, the major cytosolic HSP70, mitochondrial HSP60 and a DNAj (Fig 6D). Other mRNAs that showed less than 20% association with heat shock granules were those encoding histones, alpha and beta tubulin, 10 additional chaperones, the cytochrome oxidase complex and a few proteins involved in ribosome assembly. An ANOVA test showed that the mRNAs encoding ribosomal proteins were the only functional category that showed unique behaviour with regard to heat shock granule association (P=.00015 with Bonferoni correction). In subsequent analyses we therefore treated this group separately.

**Fig. 7.**
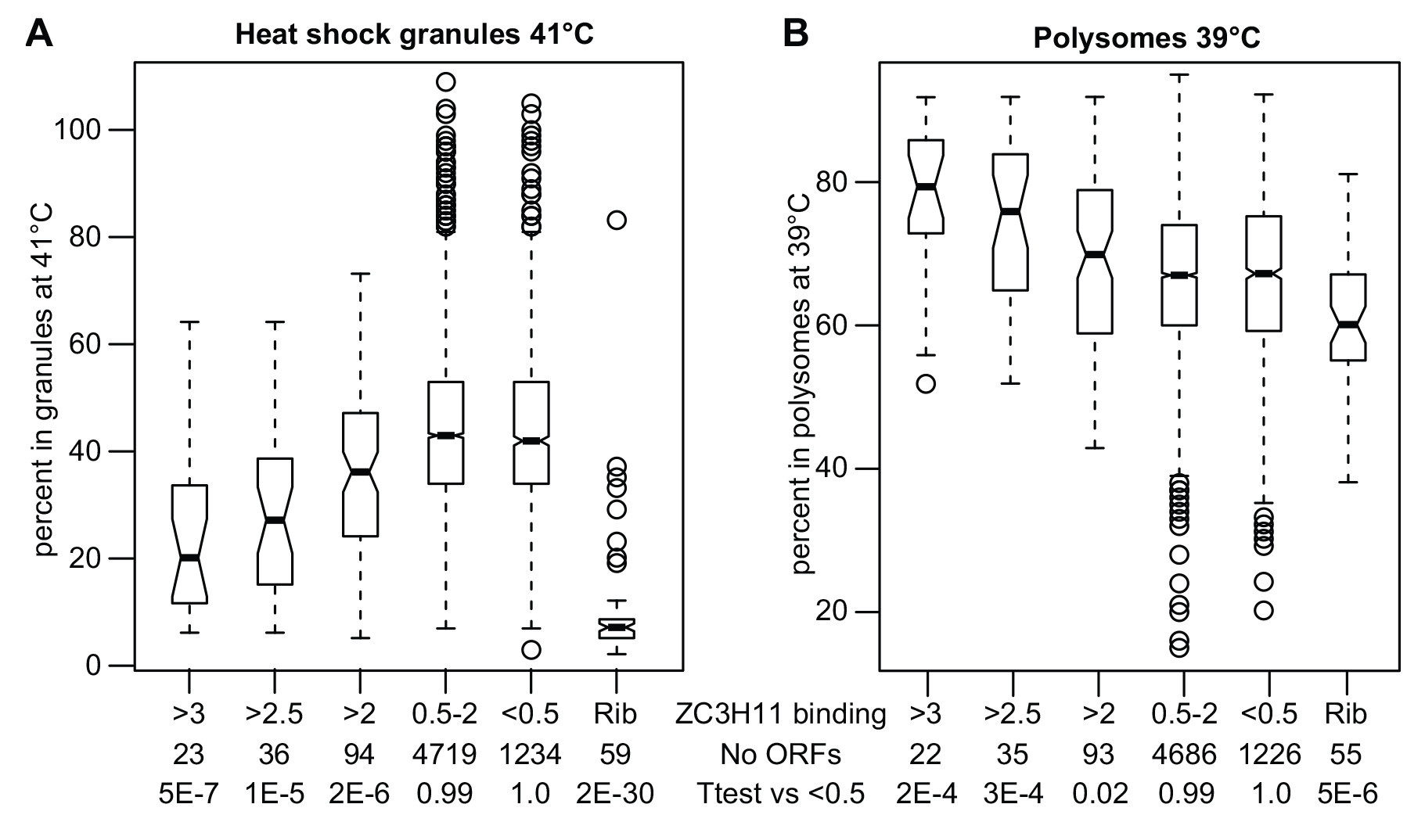
Binding to ZC3H11 related to polysome association and granule incorporation The ability of mRNAs to associate with ZC3H11 was previously assessed by co-immunoprecipitation. The extent of binding was expressed as the read count per million (RPM) in the immunoprecipitated preparation, divided by the RPM in the input. The mRNAs encoding ribosomal proteins were first extracted (Rib), then the remainder were sorted according to the bound:input ratio (indicated in the line "ZC3H11 binding"). The bottom line shows the results of a Student T-test comparing results for each category with those for mRNAs with bound:input ratios of less than 0.5. (A) Box plot showing the proportions of the mRNAs in granules (large and small combined) at41°C. (B) Box plot showing the proportions of the mRNAs in the polysomal fractions at 39°C.

After heat shock, there was little overall correlation between the coding region length and association with either total granules (S4 Fig C) or small granules alone (S4 Fig D), but DeSeq analysis showed that granules were enriched in long mRNAs (median length 4 kb) including several encoding large cytoskeletal proteins. (S5 Table, sheets 3 and 5). There was no overall correlation between loading onto polysomes at 39°C and the percentage in granules at 41°C (Fig 6E). Some potential regulators that showed reproducible mRNA abundance increases - *CYC7, DRBD5, DRBD6* and *RBP6* - showed less than 30% granule incorporation, suggesting that they might in some way be implicated in recovery from heat shock.

**Fig. 8.**
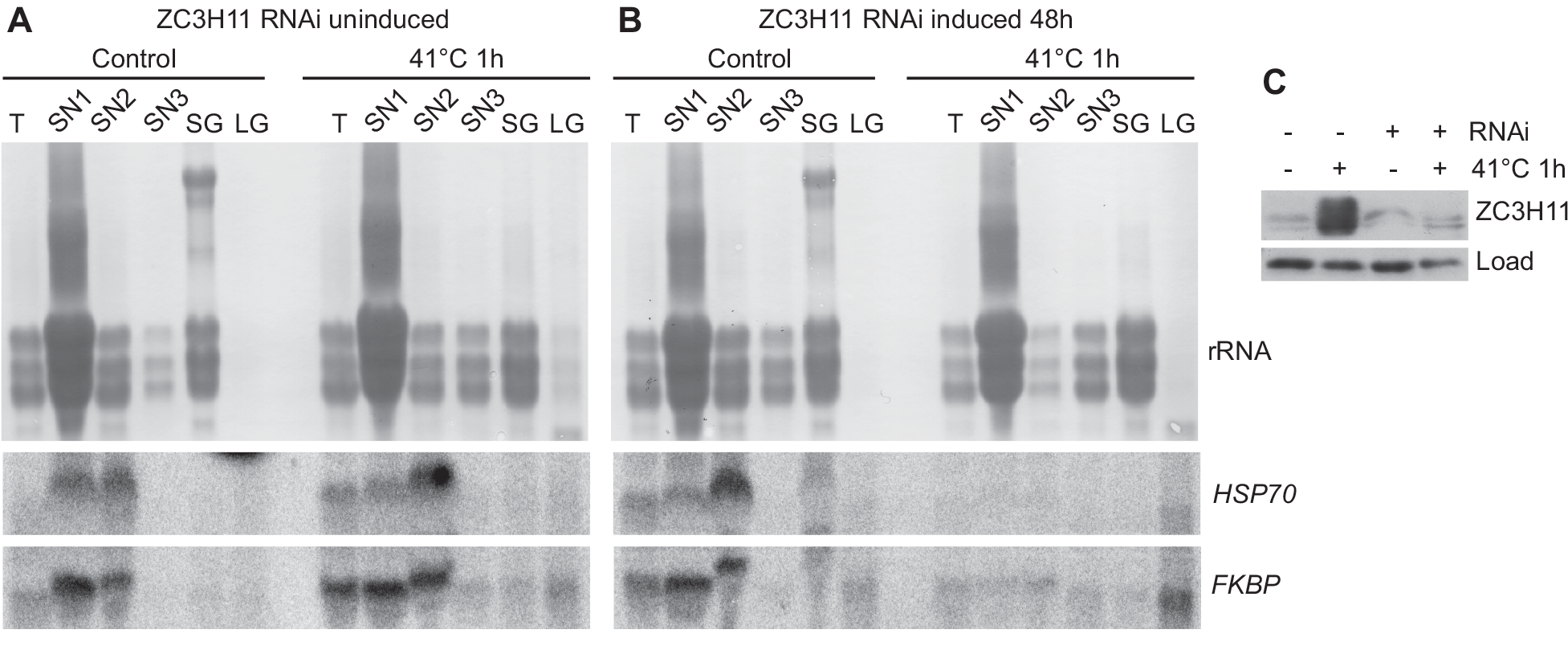
Binding to ZC3H11 protects against granule association after heat shock. A. Cells without induction of RNAi were separated into sedimentable granules (G); mRNA trapped inside cytoskeletons but not sedimented at 20000g (10min); and soluble supernatants (SN1-3). RNA was prepared from these as well as from unfractionated total cell lysate (T). All preparations are from 5*10^8^ control or heat-shocked (1 hour at 41°C) cells. RNA was analysed by Northern blotting, probing for Tb927.10.16100 (FKBP) and the major cytosolic HSP70 mRNA (Tb927.11.11330). The variations in mobility are not reproducible and might be due to different amounts of RNA and salt. B. As (A) but RNAi was induced for 48h. C. Western blot showing the amount of ZC3H11, measured as in [12]. The loading control is a cross-reacting band.

We previously showed that ZC3H11 prevents degradation of bound mRNAs after a 41°C heat shock [15]. Correspondingly, mRNAs that co-purify with ZC3H11 [15] tend to escape granule association. For the 23 mRNAs that showed the strongest enrichment in the ZC3H11-bound fraction [15], a median of 20% was associated with total granules, whereas for unbound mRNAs the median was 40% (Figs 6D and Fig 7A). A similar result was obtained if large granules alone were analysed (S4 Fig E). As previously noted, the ability to bind ZC3H11 also correlated with higher association with polysomes (Fig 7B). These results suggest that ZC3H11-bound mRNAs are protected against mRNA degradation, translational inactivation, and incorporation into granules. To check this hypothesis, we prepared granule fractions from cells with and without heat shock and / or *ZC3H11* RNAi. Without RNAi, two target mRNAs encoding HSP70 and an FKBP remained largely in the soluble fractions despite heat shock:as seen from the RNASeq results, only a tiny proportion was detected in the large granule fraction (Fig 8A). ZC3H11 depletion had no effect on this distribution without heat shock (Fig 8B, C). After heat shock, as expected, granule-free *HSP70* and *FKBP* mRNA disappeared but neither mRNA accumulated in the large granule fraction either:instead, the mRNAs were simply destroyed.

## Discussion

The results from this study have confirmed that the ability of an mRNA to bind ZC3H11 correlates not only with stablisation at high temperature, but also with continued translation and exclusion from heat shock granules. The first conclusion generalises results that were already seen for reporters with the *HSP70* 3’-UTR, while the second is consistent with previous published data indicating that mRNAs in stress granules are not translated [36–39]. The mRNAs that are bound by ZC3H11 are already quite well translated at 27°C, and become even more so after heat shock:it is possible that this high translation protects them from incorporation into heat shock granules; alternatively ZC3H11 and its associated proteins [16] might prevent granule association.

Our results show that heat shock granules are not identical to starvation granules, despite sharing some of the same proteins and mRNAs. Some mRNAs that were excluded (<20%) from heat shock granules were also similarly absent from starvation granules [20]. Presumably these encode products that are required to recover from both starvation and heat shock. Apart from ribosomal protein mRNAs, which are probably a special case and are discussed below, several chaperone mRNAs were in this category. In contrast, ZC3H11 is not implicated in the starvation response, and its mRNA was 25% in heat shock granules but 79% in starvation granules. Other mRNAs that showed a similar pattern encoded RBP3, ZC3H30, ZFP1, RBP6 and a histone H3 variant [20]. PABP1 may be important in protecting ZC3H11 target mRNAs, since it is recruited by the ZC3H11-MKT1-PBP1 complex [16]. The level of ZC3H11 protein is not increased after starvation, which explains why some ZC3H11 target mRNAs are incorporated into starvation granules (S4 Table, sheet 7).

The mRNAs encoding ribosomal proteins were almost completely excluded from both heat shock granules and starvation granules [20]. Only two annotated "ribosomal protein" mRNAs, Tb927.10.10010 and Tb927.11.6360, did not follow this pattern, but neither is a structural component of the mature ribosome. The extraordinary behaviour is therefore a universal characteristic of mRNAs that encode components of the mature ribosome. These mRNAs are also outliers in other ways:the mRNA levels are higher that would be predicted based on their half lives and gene copy numbers [4], and the average ribosome densities are relatively low (mostly less than 4 ribosomes/kb) [4,30] although the majority of the mRNAs are loaded onto polysomes [4]. Association with polysomes is also notably decreased after heat shock, without much loss of the mRNAs (Fig 2A, B):it looks as if the mRNAs are being conserved in some way other than granule sequestration. The ribosomal protein mRNAs are co-regulated during trypanosome differentiation, being decreased in stationary phase trypanosomes and increasing only 1h after addition of the differentiation stimulator cis-aconitate [40]; this is consistent with the fact that they mostly peak in the G1 phase of the cell cycle [41]. We examined the untranslated regions of these mRNAs for specific enriched motifs and found none. The only notable feature is that the 5’-UTRs are very short, with a median length of 22nt, as opposed to 108 for other mRNAs (mean±SDs are 33±34 as opposed to 203±274). Given the lack of conserved linear motifs, it is possible that secondary structures are important; or, more unusually, ribosomal protein mRNAs might be characterised by a *lack* of motifs required for recruitment of SCD6 [13] or other granule proteins. Alternatively they might be regulated via recognition of the nascent polypeptides.

Our investigation of polysome loading revealed interesting sets of mRNAs that were retained in polysomes and/or increased in abundance at 39°. Some, like ZC3H11, were rather poorly translated at the normal temperature; these migrated either near 40S, or somewhat above 40S. The reason for the difference is unknown but binding to the small subunit is unlikely [12]. The 7 mRNAs with patterns most similar to that of *ZC3H11* encoded a protein kinase, a protein phosphatase, a DNAj-like protein, and 4 other proteins of unknown function. There is no evidence of any link between these proteins and ZC3H11 function:although ZC3H11 is phosphorylated, the most likely culprit is a different kinase, casein kinase 2.1 [12]. Perhaps the most interesting observation was that the mRNAs that showed increased translation or abundance at 39°C included mRNAs that are up-regulated in salivary gland trypanosomes (Fig 2). The mammalian body environment has a temperature of 37°C (possibly higher in organs) and a 10°C temperature decrease is known to be an important factor in the switch from bloodstream to procyclic forms. However the mRNAs that increased were *not* necessarily bloodstream-form specific. Indeed, the mRNAs encoding three chaperones, two cyclins, the meiotic mRNA MND1, and the RNA-binding proteins DRBD6 and RBP6, are elevated in salivary-gland parasites but not bloodstream forms (Table 1). Induced expression of RBP6 in procyclic trypanosome cultures (at 27°C) causes differentiation to epimastigotes, and then to metacyclic forms:after 24h of RBP6 expression, about 10% of cells are epimastigotes, while metacyclics begin to appear after 5-6 days [18]. It is therefore formally possible that all of the polysomal RNA changes that we saw upon heat shock are caused by RBP6. However, this seems unlikely since the 1-h time frame is extremely short. For example, the trypanosome alternative oxidase protein appears after 2 days of RBP6 expression, but the mRNA (Tb927.10.9760) moves towards the polysomes after only an hour at 39°C (to 72% from 54%). The increased translation of *PIP39* is also intriguing. Differentiation of bloodstream forms to procyclic forms includes an intermediate called the short stumpy form. Stumpy forms are arrested in G1, and show elevated levels of PIP39. PIP39 is a protein phosphatase that is increased in stumpy forms. Further differentiation to procyclic forms is induced by addition of cis aconitate and a decrease in temperature from 37°C to 27°C, and PIP39 is essential for this transition [35]. Heat shock also increased translation of the mRNA encoding RBP7, a potential RNA-binding protein that is required for differentiation to stumpy forms [34]. It is conceivable that both PIP39 and RBP7 have functions-perhaps linked to growth arrest-in both the stumpy->procyclic and in the procyclic->epimastigote transitions. Importantly, we showed that PIP39 and RBP6 proteins also increased in procyclics incubated at 37°C, which is a temperature that is quite likely to occur in the wild.

We have absolutely no idea which stimuli initiate the development of procyclic forms to epimastigotes and metacyclic forms inside Tsetse flies. A heat shock is not required since development happens in laboratory Tsetse colonies in which temperatures are controlled below 30°C. In the field, however, Tsetse are very likely to be exposed to higher environmental temperatures, and the developing trypanosomes are exposed to warm blood meals every 3-5 days [42]. It is therefore quite possible that in the wild, temperature fluctuations inside tsetse play a role in trypanosome life-cycle progression.

## Data availability

The polysome gradient data are available under submission numbers E-MTAB-4555 and E-MTAB-4575. The heat shock granule results are available under submission number E-MTAB-4557.

## Ethics statement

No ethical approval was reqiuired for this work, which did not involve either animals or human subjects.

## Acknowledgements

We thank Chaitali Chakroborty (ZMBH) for repeating the experiment shown in Fig 8, Nicolai Kolev and Christian Tschudi for the antibody to RBP6 and Keith Matthews for the antibody to PTP1.

## Supplementary Tables

**S1 Table**

RNASeq data:effect of a 39°C heat shock analysed by DESeq. For detailed legend see Sheet 1.

**S2 Table**

RNASeq data:effect of a 29°C heat shock on the polysomal distribution of mRNAs. For detailed legend see Sheet 1.

**S3 Table**

Genes with different numbers of (AU) repeats in their 3’-UTRs.

**S4 Table**

RNASeq data:Raw results and fractions of mRNAs in heat shock granules. For detailed legend see Sheet 1.

**S5 Table**

RNASeq data:DESeq results comparing granule mRNAs with input, and total mRNA from with and without heat shock.

## Supplementary Figs

**S1 Fig.**
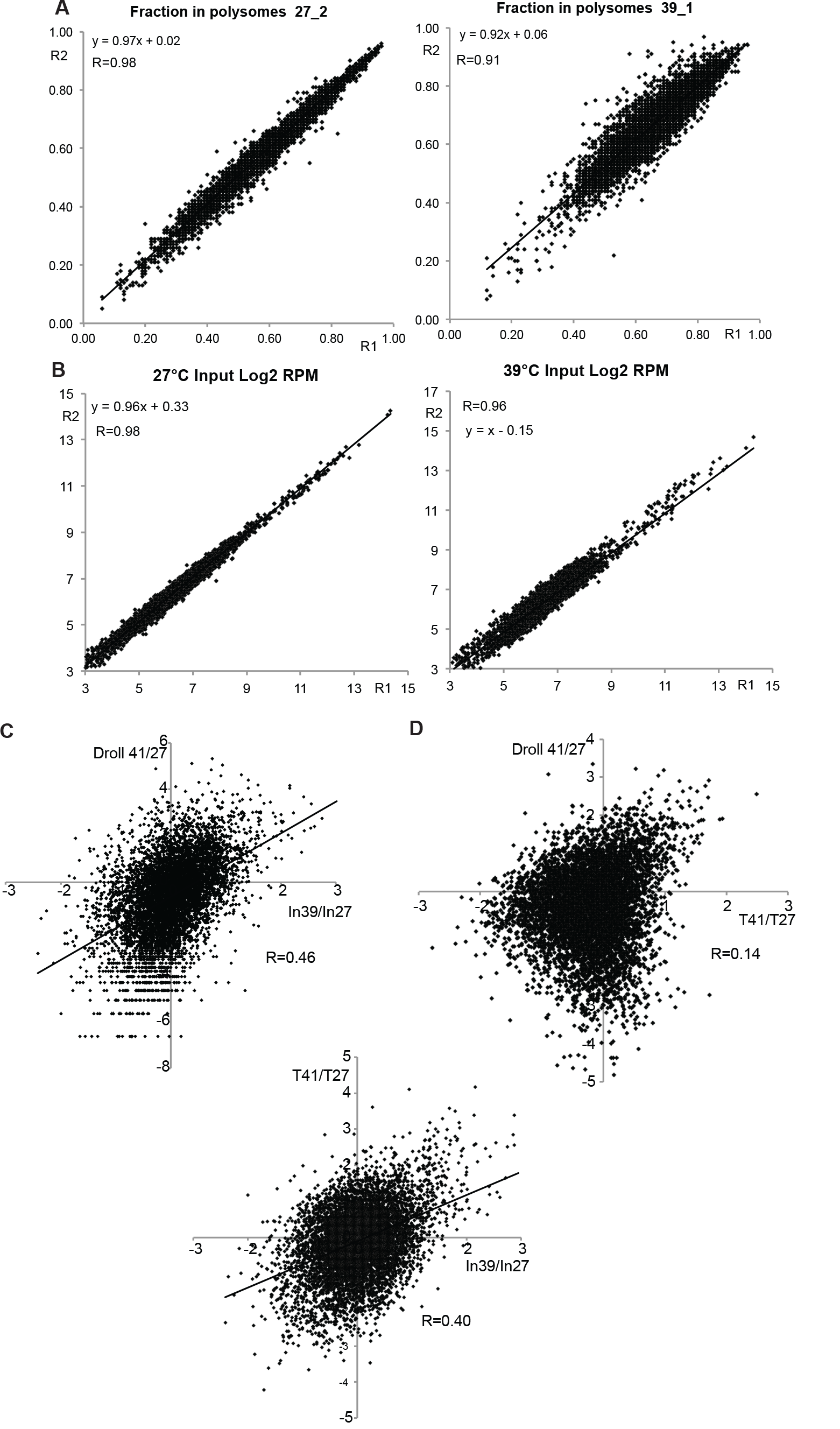
RNASeq data:correlations between replicates and comparison with previous results (A) The fractions in total polysomes at 27°C and 39°C for individual open reading frames, for replicate 1 (R1) and replicate 2 (R2). The Pearson correlation coefficient and formula for the regression line are shown. (B) As (A), but for the log2 of input reads per million (RPM). (C) Log2 of ratio of RPM values after heat shock divided by the values before heat shock.

**S2 Fig.**
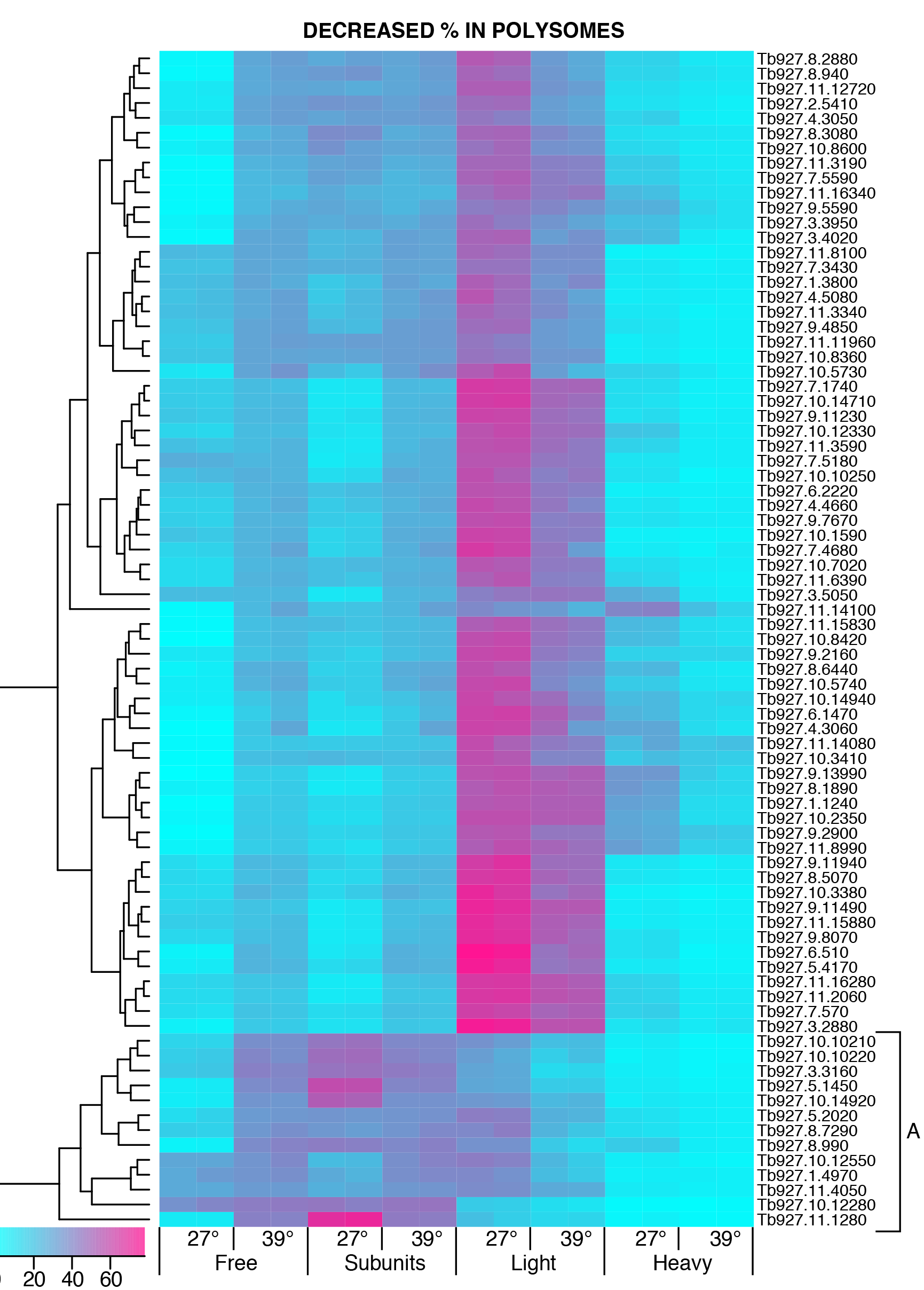
Heat map illustrating mRNAs that show reduced association with polysomes at 39°C.

**S3 Fig.**
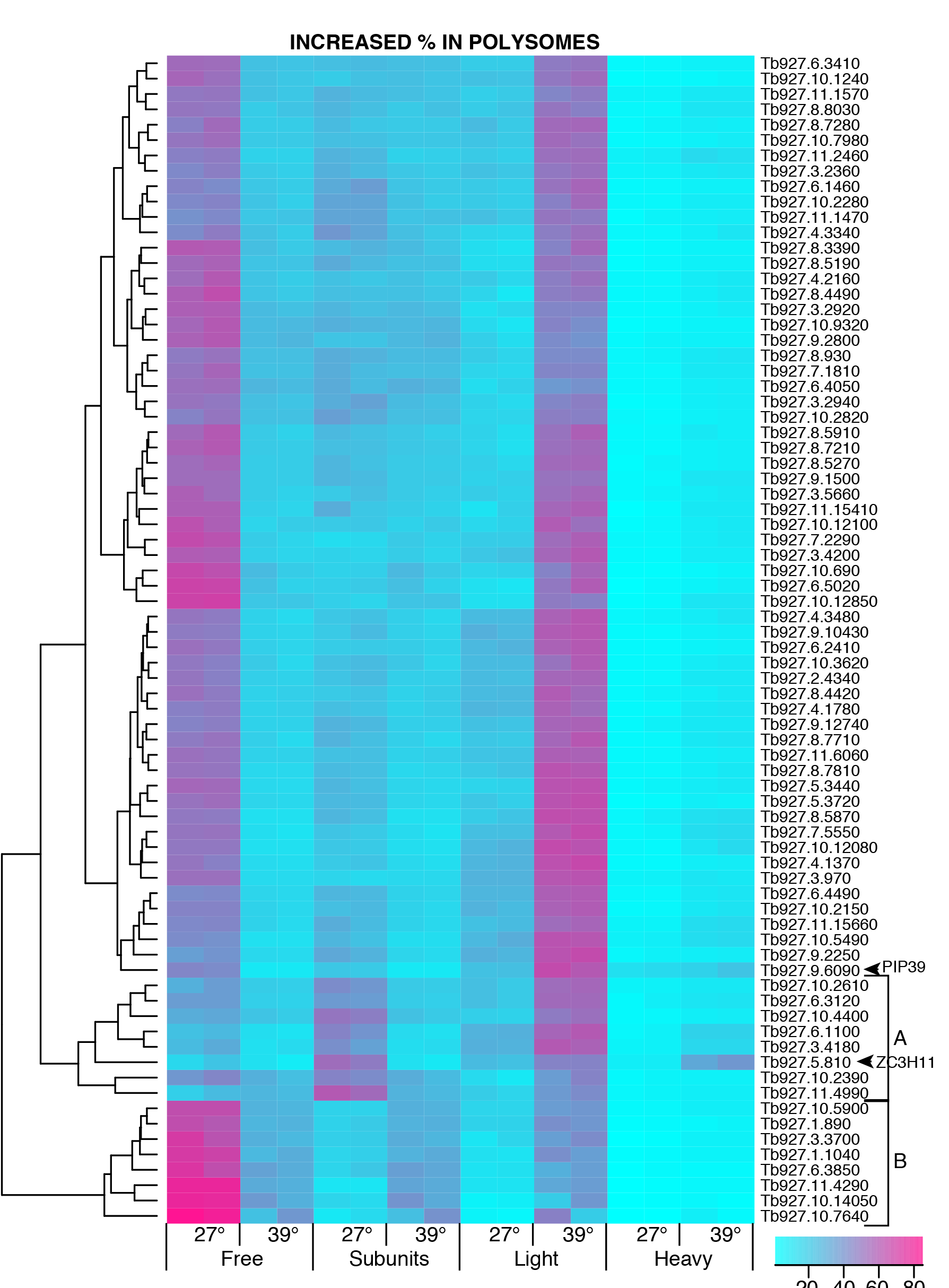
Heat map illustrating mRNAs that show increased association with polysomes at 39°C.

**S4 Fig.**
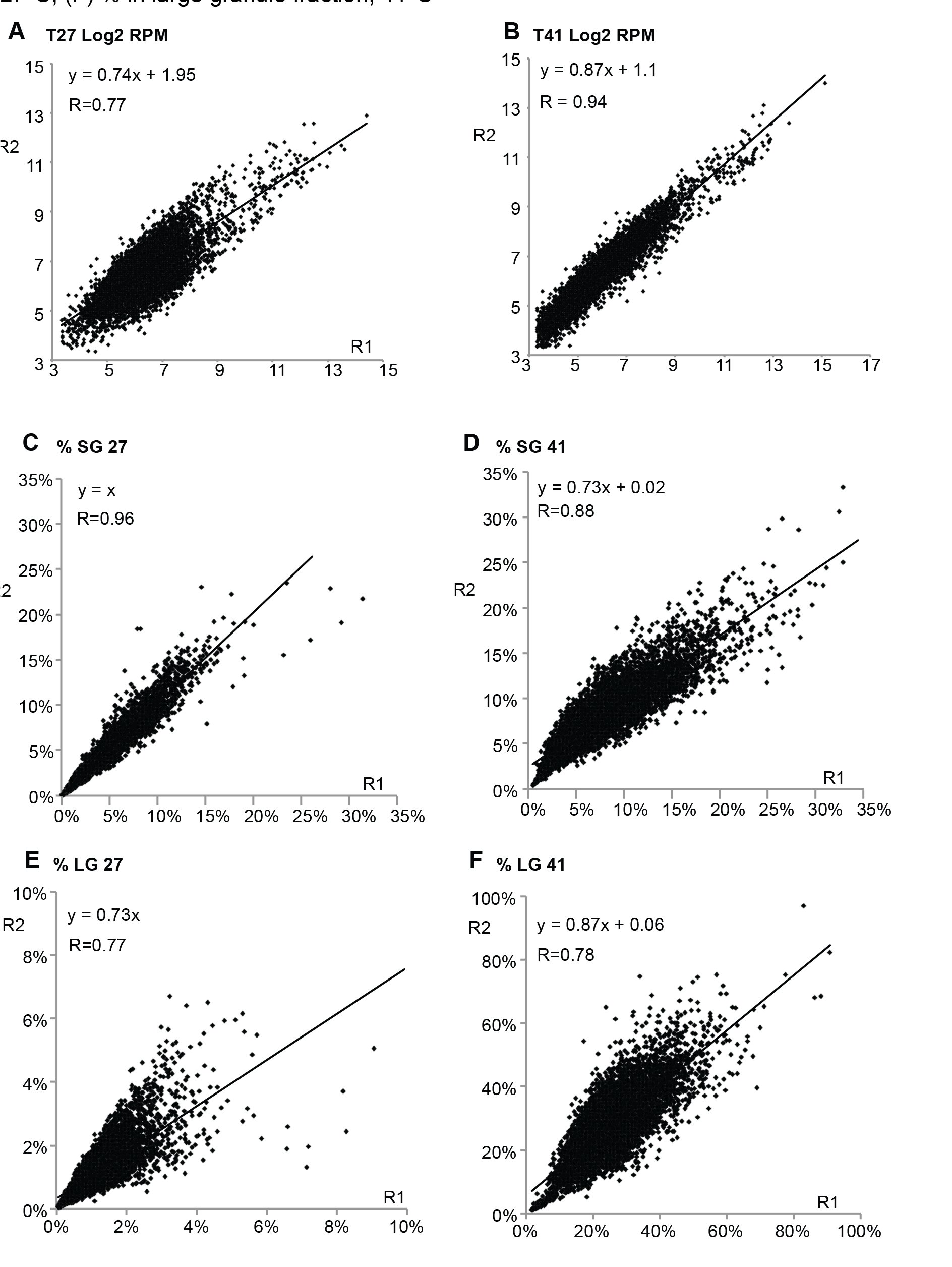
RNASeq data for granule fractionation:correlations between replicates. (A) Total RNA, 27°C-log2 of RPM; (B) Total RNA, 41°C-log2 of RPM; (C) % in small granule fraction, 27°C; (D) % in small granule fraction, 41°C; (E) % in large granule fraction, 27°C; (F) % in large granule fraction, 41°C

**S5 Fig.**
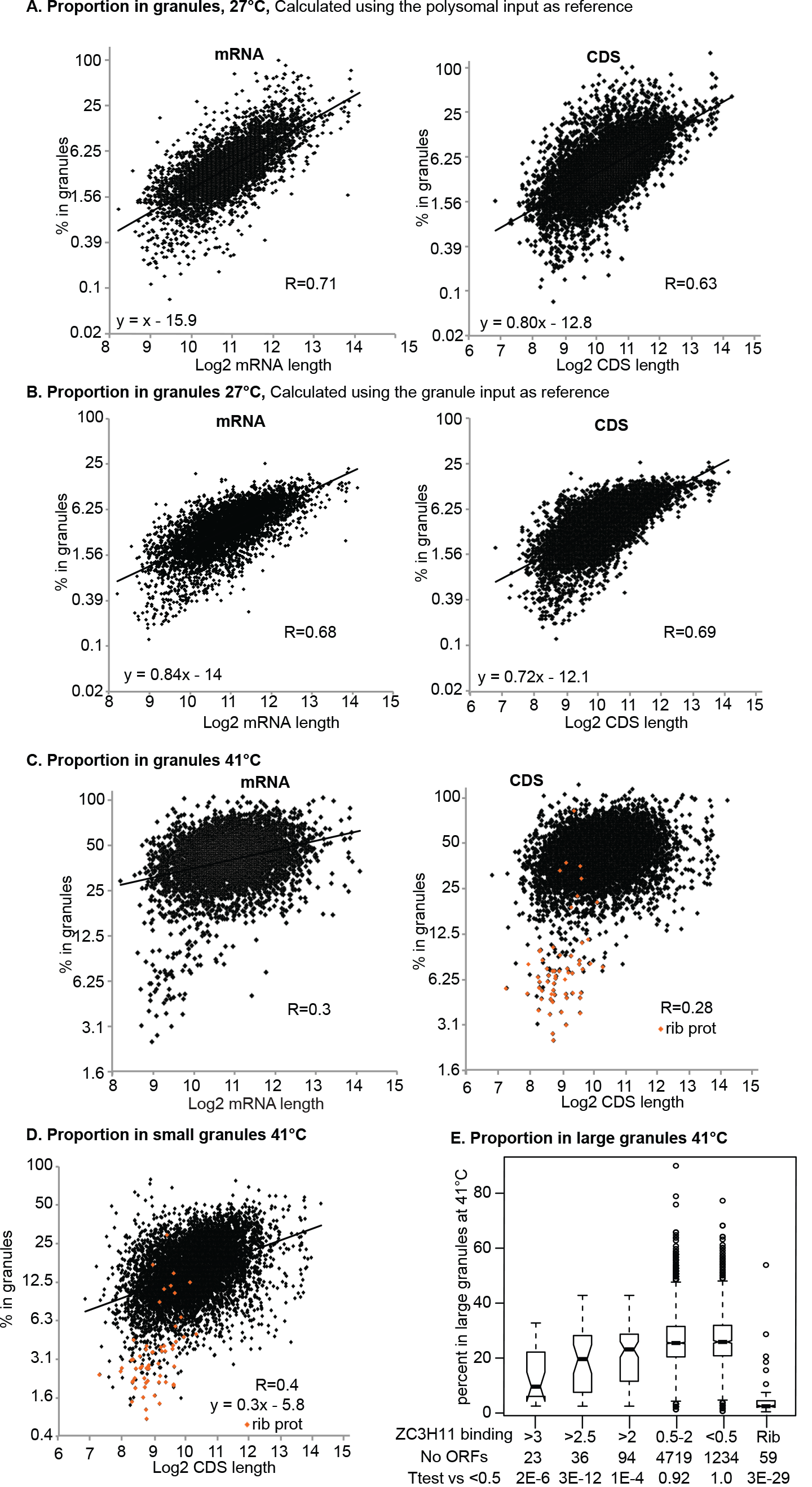
These are mostly alternative calculations to those shown in Figs 6 and 7. (A) Proportion in granules, 27°C, calculated using the polysomal input as reference instead of the total RNA, and plotted against the log2 of annotated mRNA length or coding sequence (CDS) length. (B) Proportion in granules, 27°C, calculated using the total RNA from the granule experiment as reference, and plotted against the annotated mRNA length or coding sequence (CDS) length. The left-hand panel is the same as Fig 6A. (C) Proportion in granules at 41°C plotted against the log2 of annotated mRNA or CDS length. (D) Proportion in small granules at 41°C plotted against the log2 of CDS length. (E) RNAs were grouped according to ZC3H11 binding (ratio of bound to input) and the proportion in large granules at 41°C was plotted. mRNAs encoding ribosomal proteins are shown separately.

